# Brinker regulates reciprocal outcomes of BMP signal between stem cells and differentiating cells

**DOI:** 10.1101/2025.09.14.676154

**Authors:** Samaneh Poursaeid, Jeffrey P Gamer, Mayu Inaba

## Abstract

*Drosophila* male germline stem cells (GSCs) reside at the testis tip, surrounding a cluster of niche cells known as the hub. Bone Morphogenetic Protein (BMP) ligands secreted from the hub exert both contact-dependent and -independent effects. In close proximity to the niche, BMP signaling maintains stem cells by suppressing transcription of the key differentiation factor Bag of Marbles (Bam). In contrast, the diffusible fraction of BMP promotes differentiation of cells by activating bam. How a single signaling pathway produces such opposing outcomes has remained unclear. Here, we show that the diffusible BMP fraction induces bam transcription by repressing the transcriptional repressor Brinker (Brk). We further found that *brk* mRNA displays a highly heterogeneous expression pattern within interconnected spermatogonia, suggesting that Brk may prime cell fate in a subset of transit-amplifying cells, helping to preserve a population poised for dedifferentiation while maintaining other cells for differentiation. Our findings propose a model in which a single niche-derived factor modulates reciprocal outcomes inside versus outside the niche, which is essential for the tissue homeostasis. Given the broad use of BMP signaling across stem cell niches, this mechanism may represent a general strategy to ensure correct balance between self-renewal and differentiation of stem cells.

## Introduction

Many adult stem cells divide asymmetrically to produce two daughters with distinct fates, one retains stem-cell identity, while the other initiates differentiation. Although asymmetric division is an efficient strategy for stem-cell maintenance, stem cells have finite lifespans, and lost cells can be replenished either through symmetric self-renewal divisions or by dedifferentiation^1-6^. We previously demonstrated that dedifferentiation, where partially differentiated cells revert to stem-cell identity, is the predominant mechanism for maintaining germline stem cells (GSCs) in the *Drosophila* testicular niche ^1,7,8^. Despite its importance, the molecular and cellular mechanisms governing dedifferentiation remain poorly understood.

The *Drosophila* testis provides a simple and powerful model to study stem-cell regulation in the niche. Each testis harbors 8–10 GSCs that directly contact a central cluster of niche cells, the hub (Figure 1A)^9,10^. Upon GSC division, one daughter remains attached to the hub and retains stem-cell identity, while the other initiates differentiation program, four synchronous transit-amplifying divisions to form a 16-cell spermatogonia (SGs) (Figure 1A)^4,11-14^. The Bone Morphogenetic Protein (BMP) ligands, Decapentaplegic (Dpp) and Glass bottom boat (Gbb) are secreted from the hub in addition to a cytokine-like ligand, Unpaired (Upd). These ligands are essential for GSC maintenance^15-18^. In GSCs, BMP ligands strongly activate its canonical pathway and phosphorylated Mad (pMad) suppress transcription of the key differentiation factor Bam^16-20^. When a daughter cell loses hub contact, bam expression is activated, allowing entry into the differentiation program^16-20^.

**Figure 1.**
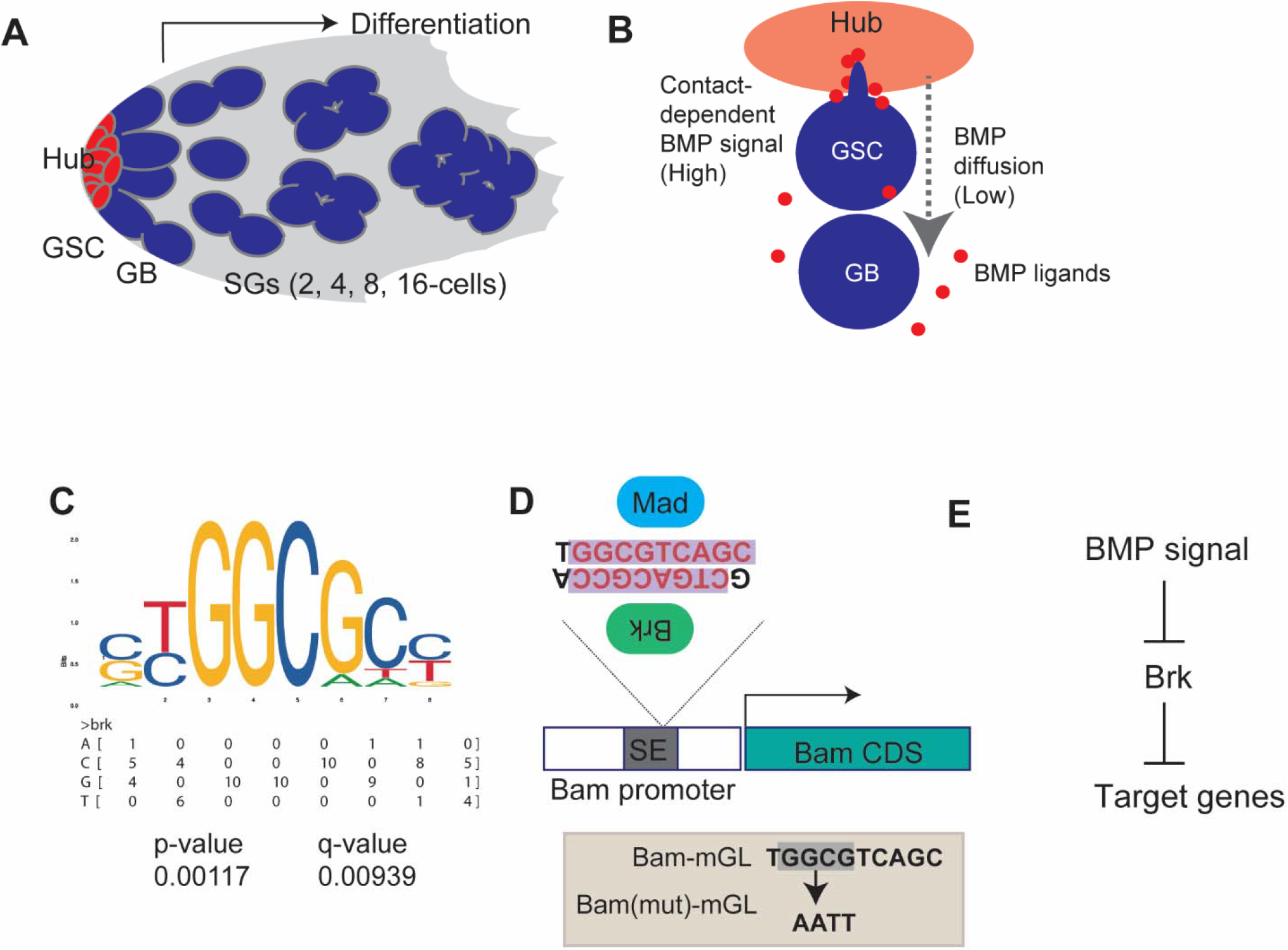
Bam promoter contains putative Brinker (Brk) binding site. (A) The diagram shows anatomy of *Drosophila* testicular niche. GSC: Germline stem cell, GB: Gonialblast, SGs: Spermatogonia. (B) Asymmetric division of a male GSC. BMP signal mediates distinct outcomes between GSC and GB/SG populations via contact dependent and independent mechanisms, respectively. GSC possesses specialized cellular protrusion, called microtubule based nanotubes that enhance the signal reception specifically in GSCs. (C) Brk binding motif obtained from JASPER (matrix profile; MA0213.1) and nucleotide frequency matrix. P-value and q-value were obtained for bam promoter sequence by motif scanning. (D) Structure of bam promoter. Previously identified Mad binding sites are potentially recognized by Brk on its antisense strand. Lower box shows mutated nucleotides for Bam(mut)-mGL reporter construct. All reporters contain bam promoter from position −198 relative to TSS to endogenous start codon of bam gene. (E) Suggested relationships between BMP signal and Brk.

In addition to hub cells, Gbb is also expressed in somatic cyst cells that encapsulate differentiating SGs and regulates proper differentiation of SGs^21^. In contrast, Dpp is exclusively expressed in the hub^22^, and was thought to act strictly in a short range, restricted to GSCs in direct hub contact through specialized signaling protrusion, called microtubule-nanotubes, ensuring that signaling is limited to stem cells and excluded from differentiating daughters^23,24^. However, our previous work revealed that a diffusible fraction of Dpp does exist outside the niche and exerts the opposite effect on bam expression. While niche-bound Dpp represses *bam* expression in GSCs, diffusible-fraction of Dpp promotes *bam* expression in differentiating cells, thereby preventing excessive dedifferentiation^22^. These opposing outcomes are likely determined by different strength of the signal, that creates distinct concentration of Mothers against Dpp (Mad), the downstream transcription factor of BMP pathway (Figure 1B)^22^. However, the molecular basis for this dual action remains unclear.

Brinker (Brk), a transcriptional repressor that antagonizes BMP signaling, plays a critical role in multiple developmental contexts. BMP signaling directedly suppresses brk transcription, establishing a reciprocal relationship between Dpp activity and Brk expression^25-28^. This antagonistic pattern allows Dpp to indirectly activate specific target genes by repressing *brk* transcription. Such toggling is essential for precise gene regulation, supporting proper tissue patterning and cell differentiation in response to developmental cues, as observed in various systems including the embryo ^26,28^, wing imaginal disc ^25,27^, midgut ^28^, and egg chamber ^29^. Given that Dpp and Brk exhibit a reciprocal regulatory relationship in other tissues, we anticipate that a comparable mechanism may exist in the testis.

In this study, we investigate the role of Brk regulation of *bam* expression in the testis. Our findings suggest that Brk is expressed in differentiating cells and mediates the opposing outcome of BMP signaling from GSCs.

## Results

### Bam promoter contains putative Brinker (Brk) binding site

In *Drosophila* germline stem cells (GSCs), bam transcription is repressed by niche-derived BMP signaling^30,31^. This repression occurs through a silencer element (SE) within the bam promoter, which is directly bound by the canonical BMP pathway effector, Mad and its partner Medea (Med)^30^.

Our previous study has demonstrated that BMP signaling modulates *bam* transcription through the SE in a location-dependent manner, such that suppressing *bam* transcription in GSCs while enhancing it in differentiating SGs^22^. This dual activity raises the possibility that multiple transcriptional regulators act on the bam promoter, and that the interplay between these factors downstream of BMP signaling determines the final transcriptional outcome.

A strong candidate for such a regulator is Brk, a transcriptional repressor that is a well-established antagonist of BMP signaling across diverse developmental contexts^26,32^. Brk typically binds to the promoters of BMP-responsive genes to repress their transcription, while BMP signaling itself represses *brk* expression, thereby allowing target gene activation^32^.

To assess the possibility that Brk directly influences *bam* transcription, we performed motif-scanning analysis of the SE using FIMO^32-34^ with the Brk consensus binding sequence obtained from JASPAR^34^. Strikingly, this analysis revealed that the previously determined Mad-binding site within the SE overlaps with a potential Brk-binding site (Figure 1C, D)^31^. This observation suggested a unique relationship between Mad and Brk at the SE, in which BMP signaling might indirectly activate *bam* expression by suppressing *brk* expression, as documented in other systems (Figure 1E)^35^.

### Brk suppresses *bam* transcription in SGs

Given the presence of a putative Brk-binding site within the bam promoter, we next asked whether Brk directly represses *bam* expression in SGs. To study *bam* transcriptional regulation, we used a previously established transcriptional reporter, in which the *bam* core promoter (198-bp upstream of the transcription start site) drives expression of the fluorescent protein mGreenLantern (Bam-mGL)^22^. Consistent with endogenous *bam* expression, Bam-mGL was strongly activated beginning in the 4-cell SGs, and remains high in later SGs (Figure 2A)^22^. To test the role of Brk in *bam* expression, we examined Bam-mGL activity under conditions of perturbation of Brk levels specifically in SGs. Brk knockdown under the BamGal4 driver (Bam>Brk RNAi) led to a marked increase in Bam-mGL intensity in SGs, whereas overexpression of Brk (Bam>Brk) caused a significant reduction of Bam-mGL signal in both 4-cell and 8–16 cell SGs compared with controls (Figure 2A–D). Importantly, the effect of Brk RNAi was abolished when the Bam(mut)-mGL reporter was used, in which the putative Brk-binding site was disrupted (Figure 2E–H). These results strongly suggest that Brk directly binds to the SE in the *bam* promoter to regulate reporter activity.

**Figure 2.**
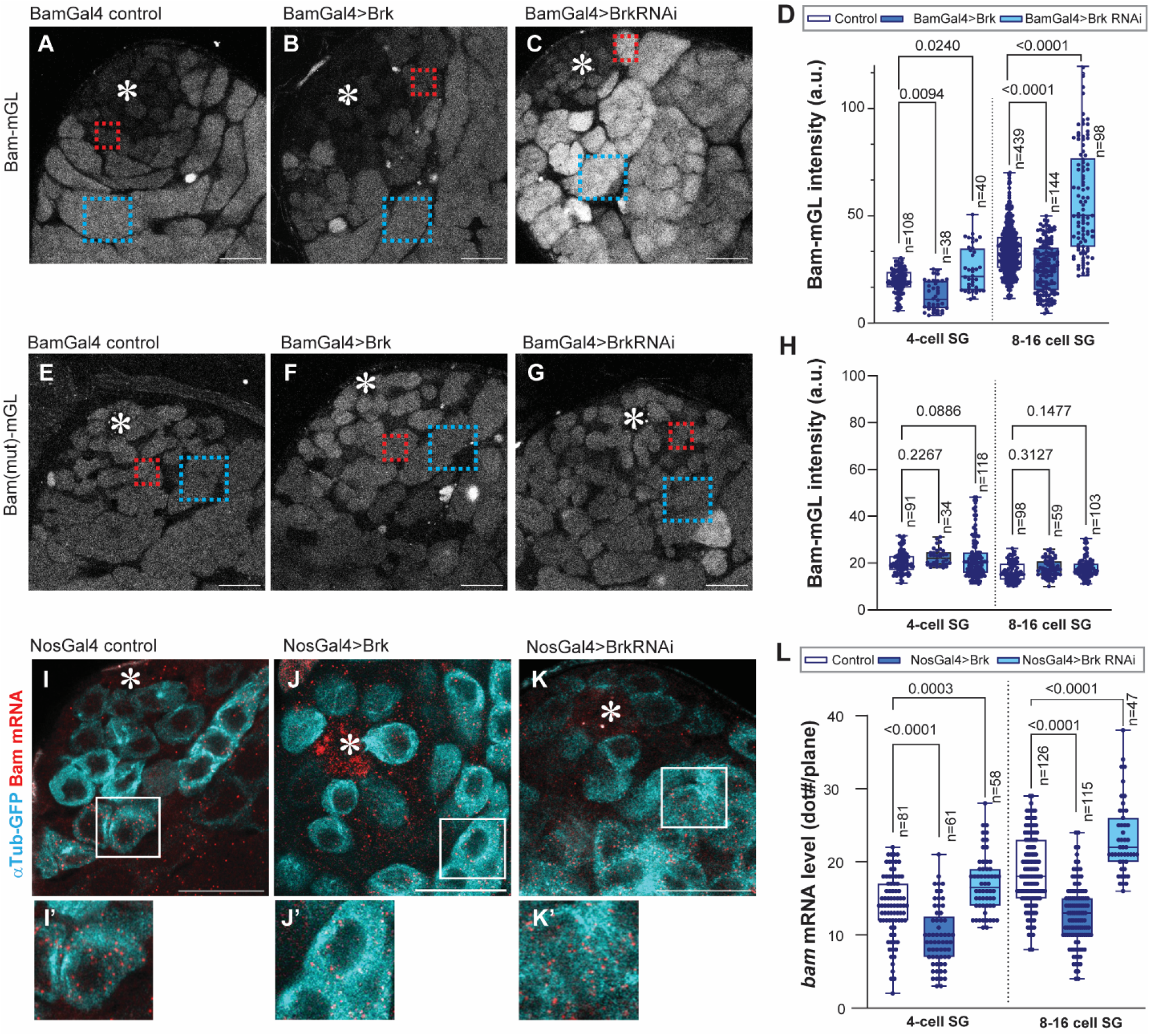
Brk suppresses bam transcription in SGs. (A-C) Representative images of Bam transcriptional reporter (Bam-mGL) in indicated genotypes. (D) Intensity quantification of Bam-mGL reporter in 4- or 8-16 cell SGs for indicated genotypes. n indicates the number of scored cysts. (E-G) Representative images of mutated version of Bam transcriptional reporter, Bam(mut)-mGL, in indicated genotypes. (H) Intensity quantification of Bam(mut)-mGL reporter in 4- or 8-16 cell SGs for indicated genotypes. n indicates the number of scored cysts. Squared regions in the images (A-C, E-G) indicate examples of measured area on 4-cell SG stage (red) or 8-16-cell SG stage (blue). (I-K) Testis-tip images of bam mRNA FISH (red) for indicated genotypes. Germ cells are visualized by NosGal4>αTub-GFP (Cyan). Lower panels (I’-K’) show magnified images of squared area in I-K. (L) mRNA quantification from mid-plane of cells in indicated stages. mRNA molecules (smFISH dots/plane) were manually counted in the cells in indicated stages. n indicates the number of scored cells. All box plots show 25–75% (box), median (band inside) and minimum to maximum (whiskers) with all data points. For Brk overexpression, UAS-Brk (#90380) was used. p-values were calculated by Šídák’s multiple comparisons tests in all graphs and provided on each graph. Asterisks indicate the hub. All scale bars indicate 20µm.

To validate these findings at the endogenous locus, we performed single-molecule fluorescent in situ hybridization (smFISH) against *bam* mRNA^36^. Consistent with the reporter assays, Brk knockdown significantly increased the number of *bam* mRNA molecules in SGs, whereas Brk overexpression led to a substantial reduction in *bam* transcripts (Figure 2I–L). Together, these results indicate that Brk suppresses excessive *bam* transcription in SGs by directly acting on the SE, located within the *bam* promoter.

### BMP signal suppresses Brk expression in SGs

In early embryogenesis and wing imaginal disc development, Brk represses BMP-responsive genes, and its own transcription is negatively regulated by Dpp signaling through the Mad-Medea-Schnurri complex^37^. Similarly, in the *Drosophila* ovary, the expression of *brk* is suppressed by BMP signal^29^. Therefore, we sought to test whether the same regulation is also utilized in the testis.

We first examined the expression pattern of Brk in the testis by in situ hybridization chain reaction (HCR) using previously validated probe against *brk* mRNA^38^. We detected *brk* mRNA in both germline and somatic cells (Figure 3A). In germline, *brk* mRNA is absent in GSCs and mainly observed in 4- to 8-cell SGs, the stage at which bam expression is positive^39^ (Figure 3A).

**Figure 3.**
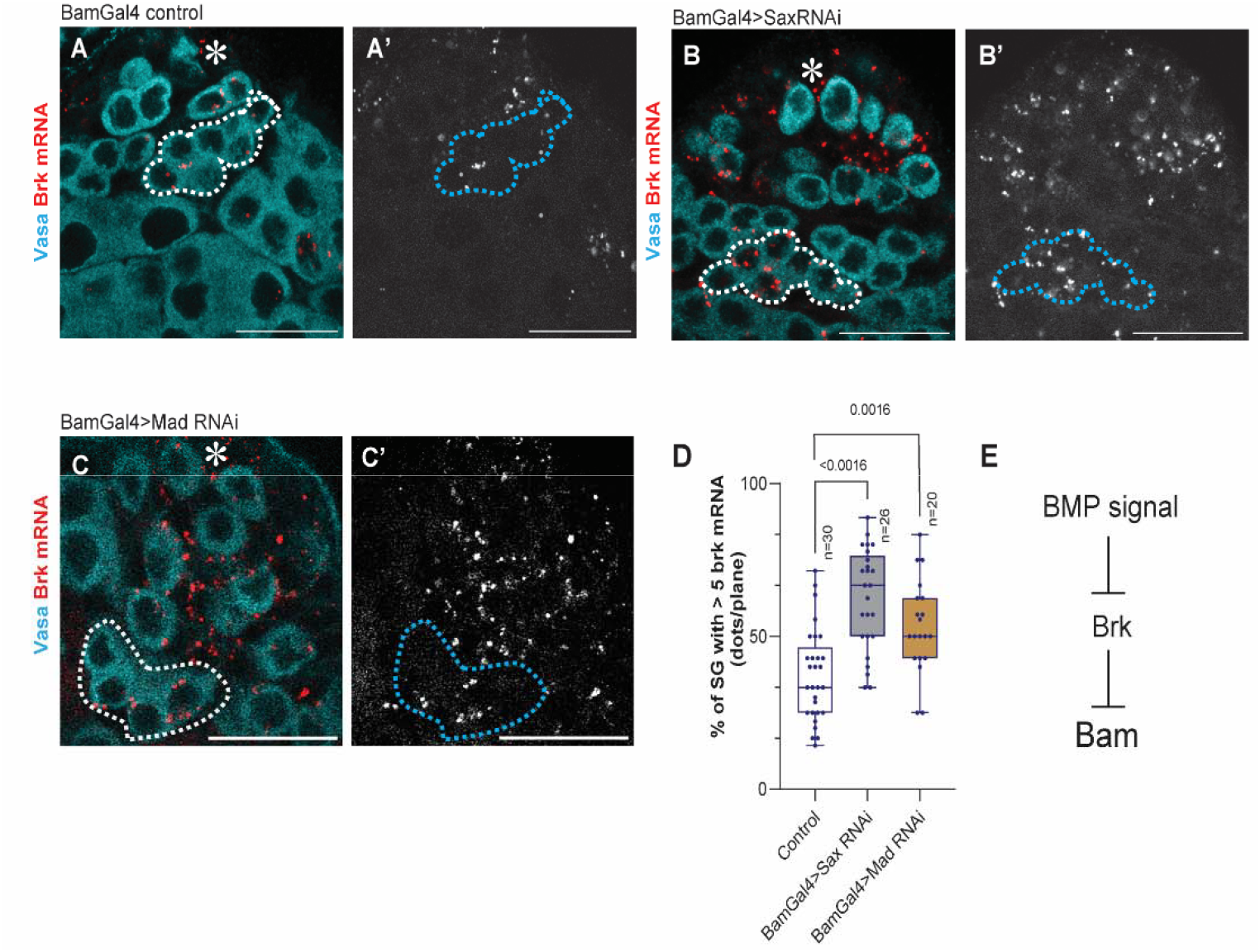
BMP signal suppresses Brk expression in SGs. (A-C) Representative testis-tip images of brk mRNA (red) for indicated genotypes. Germ cells are visualized by Vasa staining (Cyan). Broken lines encircle 8-cell SG cysts. (D) The graph shows frequency of the SG cysts with brk-high cells (containing more than 5 brk mRNA). HCR dots were counted from mid-plane cells in SGs. n indicates the number of scored testis. Box plots show 25–75% (box), median (band inside) and minimum to maximum (whiskers) with all data points. Note that distributions of each SG stage were similar across examined genotypes, indicating that the observed change is not due to SG differentiation defect. p-values were calculated by student-t-test. (E) Suggested model based on the results. Asterisks indicate the hub. All scale bars indicate 20µm.

To test whether BMP signaling represses *brk* expression in SGs, we knocked down the type I receptor Sax, the primary mediator of diffusible BMP signal in SGs ^22^, and Mad, the downstream effector of BMP signaling. Knockdown of either Sax or Mad under the BamGal4 driver led to a significant increase in the number of *brk* transcripts in SGs (Figure 3A-D), indicating that BMP signaling negatively regulates *brk* transcription in SGs (Figure 3E).

### Brk is required for maintenance of stem cell pool

In the *Drosophila* testis, GSCs are primarily maintained through asymmetric division, producing one self-renewing GSC and one differentiating daughter cell^11^. However, GSC loss occurs at a certain frequency. Therefore, additional mechanisms are required to sustain GSC population. Our previous study demonstrated that dedifferentiation, whereby displaced daughter cells revert to a GSC identity by re-entering the niche, plays a significant role in maintaining the GSC population^7,40^.

Our previous study has demonstrated that diffusible fraction of Dpp, a major BMP ligand in the niche, promotes SG differentiation by suppressing excess dedifferentiation through activation of *bam* expression^22^. Since Brk represses *bam* expression in SGs, we hypothesized that Brk may be a key factor that induce dedifferentiation via downregulating *bam* transcription. To test this possibility, we examined whether perturbing Brk levels in the germline affects long-term GSC maintenance or not.

Knocking down of Brk under the BamGal4 or NosGal4 driver both resulted in a gradual reduction in GSC number over the fly age, reaching significant by 21 days post-eclosion compared to the control (Figure 4A, B, Figure S1). These effects are contrary to the Sax knock down phenotype which results in gradual increase of GSC number shown in previous study^22^, suggesting that Brk removal likely blocks dedifferentiation.

**Figure 4.**
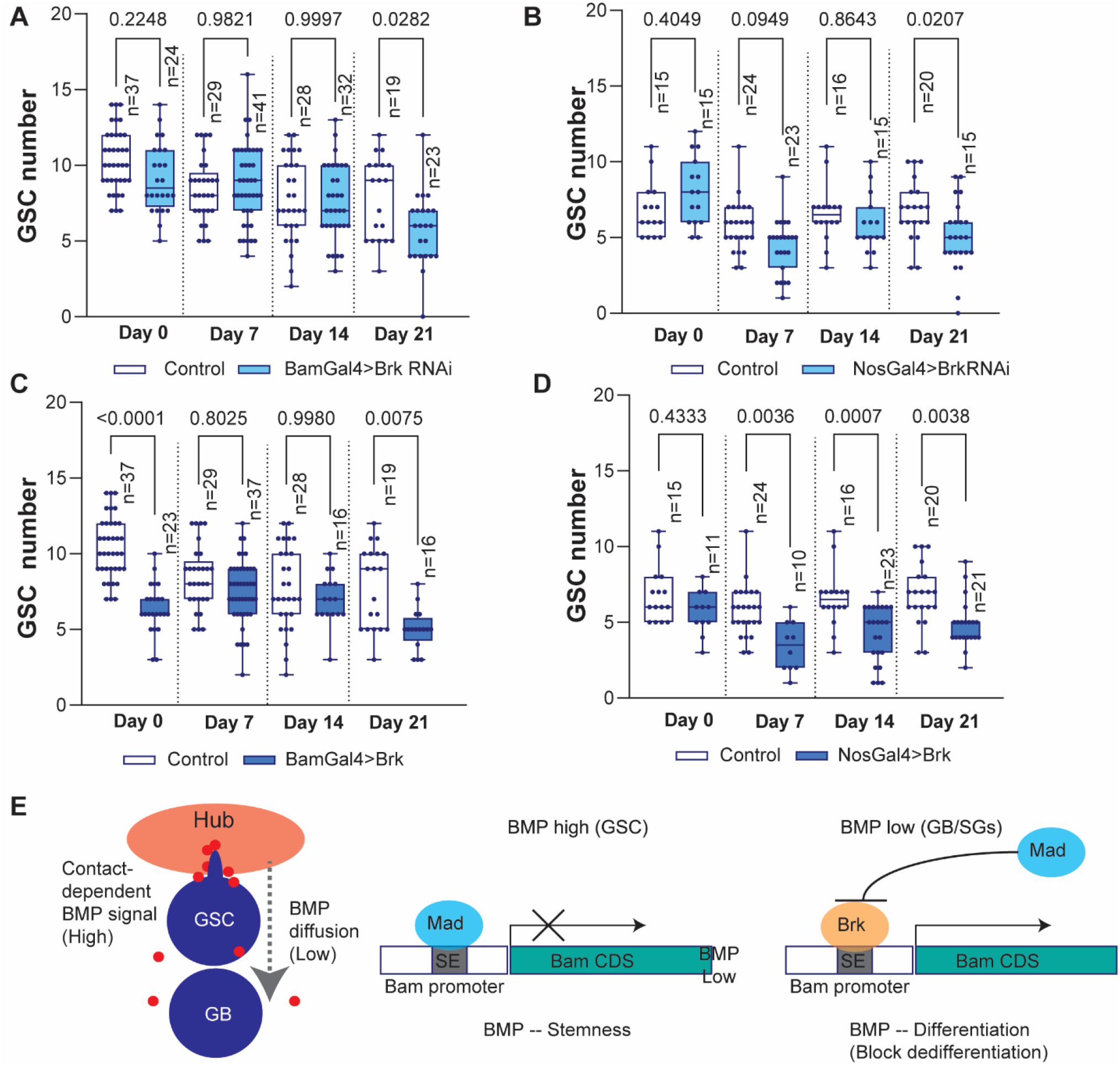
Brk is required for maintenance of stem cell pool. (A-D) Changes in GSC numbers during aging in the testes isolated from indicated genotypes. p-values were calculated by Šídák’s multiple comparisons tests in all graphs and provided on each graph. n indicates the number of scored niches (testes). GSC number was scored by confocal imaging of testis tips (see Figure S1 for example). (E) Suggested model based on the results. All box plots show 25–75% (box), median (band inside) and minimum to maximum (whiskers) with all data points.

Next, we examined the effect of Brk overexpression. If Brk is an essential factor that induces dedifferentiation, overexpression Brk may accelerate the process and lead to gradual increase of GSC number. However, unexpectedly, we found that overexpression of Brk caused a significant decrease in GSC number throughout all ages via an unknown effect, making it difficult to judge its effect on dedifferentiation (Figure 4C, D). The effect of Brk overexpression on low GSC number is already present in the young flies (day0-7 post-eclosion), suggesting that Brk may play a role in determination GSC number during the niche establishment process independent of its effect on dedifferentiation.

Taken together, these data suggest that the Brk is an essential factor for maintaining the GSC pool in the niche via regulation of *bam* transcription. The SE integrates both positive and negative regulatory inputs downstream of BMP signaling, with Mad and Brk acting as repressors that bind overlapping site in a stage-specific manner. Mad is phosphorylated and activated by BMP signaling, whereas *brk* expression is suppressed by the same pathway (Figure 4E). This opposing effect of the BMP signaling on distinct binding factors may determine whether *bam* remains repressed in GSCs or becomes upregulated in differentiating progeny.

### Brk gene product displays heterogenous distribution in SG cysts

While examining *brk* expression, we observed notable heterogeneity in transcriptional levels of *brk* among cells within individual SG cysts (Figure 5A). In 2–16-cell cysts, *brk* mRNA was predominantly enriched in a small subset of cells, whereas others displayed little or undetectable levels. To determine whether this heterogeneity arises from transcriptional variation or post-transcriptional regulation, we analyzed the distribution of transcripts from a UAS-HA-Brk transgene. As seen in endogenous *brk*, transgene-derived mRNA accumulated variably in only a small fraction of SG cells, correlating with protein levels detected by HA staining (Figure 5B–C), suggesting that the heterogeneity is subject to post-transcriptional regulation. Consistent with this, ectopic Dpp expression, which causes an accumulation of SG-like cells and a tumor-like phenotype, *brk* mRNA remained heterogeneously distributed among SGs (Figure 5D). This indicates that heterogenous localization of *brk* mRNA occurs within interconnected SG cysts. To test whether this phenomenon is specific to Brk, we examined UAS-HA-Mad expression. In contrast to Brk, HA-Mad was detected uniformly across all cells within SG cysts (Figure 5E), suggesting that the variability is a unique feature of the *brk* gene.

**Figure 5.**
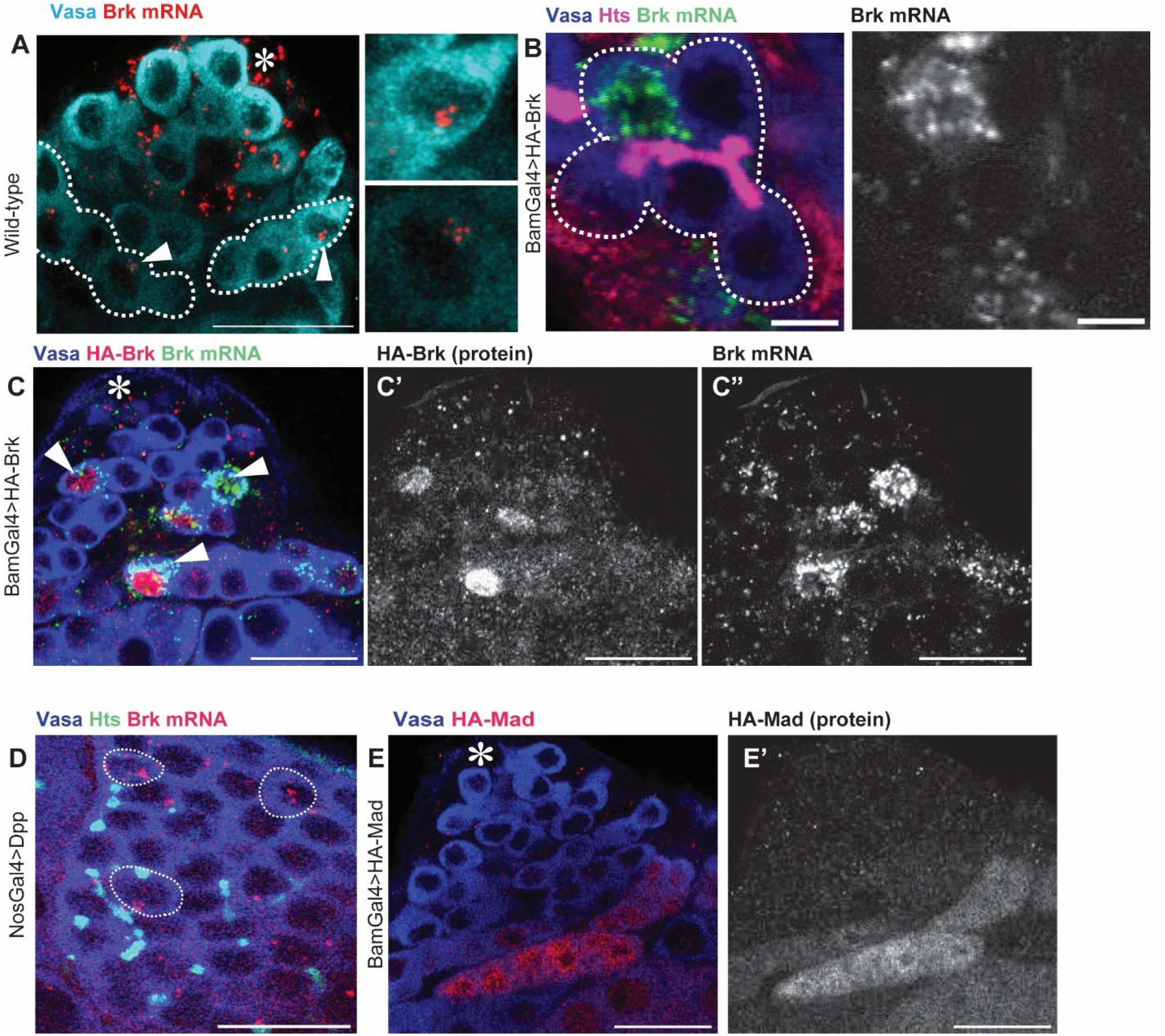
Brk gene product displays heterogenous distribution in SG cysts. (A) Representative testis-tip images of *brk* mRNA (red) in wild type (yw). Germ cells are visualized by Vasa staining (Cyan). Broken lines encircle 8-cell SG cysts. Arrowheads indicate clustered *brk* mRNA in cells. (B) An example of a 8-cell SG cyst showing biased distribution of overexpressed *brk* mRNA (BamGal4>HA-Brk). Connection of germ cells is visualized by fusome (Hts staining, red). The broken line encircles a cyst. Right panel shows a *brk* mRNA channel. (C) Representative testis-tip images of *brk* mRNA (red) and HA staining in Brk overexpressing testis (BamGal4>HA-Brk). (D) Representative image of *brk* mRNA (red) in UAS-Dpp. Germ cells are visualized by Vasa staining (blue). Arrowheads indicate Brk highly expressed cells within SG cysts. (E) Representative testis-tip images of HA staining in Mad overexpressing testis (BamGal4>HA-Mad). Germ cells are visualized by Vasa staining (blue). Asterisks indicate the hub. Scale bars indicate 5µm for B, 20µm for other images.

## Discussion

Using *Drosophila* male gonads, we previously showed that diffusible Dpp plays a critical role outside the niche by preventing excess dedifferention of GSC daughter cells, a function opposite to its niche role of promoting GSC self-renewal. Remarkably, these contrasting outcomes are mediated through the same canonical BMP pathway, which represses *bam* expression in stem cells but upregulates *bam* in differentiating cells^22^. In this study, we found that the transcriptional repressor Brk plays a role on this location-dependent switch in *bam* regulation. Brk is specifically expressed in *bam*-positive populations, where it lowers *bam* transcript levels. At the same time, BMP signaling outside the niche represses *brk* expression, thereby relieving repression and enhancing *bam* expression.

Brk is a key component of the BMP signaling pathway, functioning as a transcriptional repressor for a broad range of BMP target genes^25-28^. Typically, Brk expression displays a pattern reciprocal to pMad levels^25-28^. In the testis, BMP signaling is strongly activated near the hub, as indicated by high pMad staining in GSCs, where pMad suppresses *bam* transcription directly. A diffusible fraction of Dpp secreted from the hub travels to the SGs, weakly activating BMP signaling, where the activation is essential for increasing *bam* expression^22^. Interestingly, Brk is specifically expressed in SGs, where BMP signaling remains active^22^. This raises the intriguing possibility that the *brk* promoter may exhibit higher sensitivity to pMad than the *bam* promoter. Further studies will be needed to determine the relative pMad-binding affinities in vitro to these two promoters.

Intriguingly, we identified remarkable heterogeneity in Brk expression level within SG cysts. It has been widely assumed that cells within *Drosophila* SG cysts, which are physically interconnected by fusomes, are equivalent and contribute equally to sperm formation^41^. However, our findings revealed, for the first time, an example of molecular heterogeneity within these syncytial cysts, despite their cytoplasmic continuity. Bam expression is known to distribute equally in interconnected SGs^20^. Prior to the dedifferentiation, SG cysts are fragmented and migrate back to the niche^2^. Further experiments are needed to determine which genes are influenced by Brk heterogeneity and how Brk-positive subpopulations behave.

In summary, our findings indicate that Brk is a critical integrator of diffusible niche-derived Dpp signals controlling *bam* expression in SGs. By acting as a repressor at the *bam* promoter, Brk ensures that *bam* is expressed at appropriate levels to prevent excess dedifferentiation. Whereas it creates a highly heterogenous population within interconnected SG cysts likely to secure cells primed for dedifferentiation. Our work provides the mechanism in which a single niche ligand induces distinct cellular responses inside versus outside the niche, which may be a common mechanism to regulate tissue homeostasis.

## Materials and Methods

### Fly husbandry and strains

Flies were raised on standard Bloomington medium at 25°C, unless temperature control was required. Adult flies (0- to 7-days old) were used in all experiments. The following fly stocks were sourced from the Bloomington Drosophila Stock Center (BDSC): UAS-Brk (BDSC90380); brk RNAi (TRiP.HMC03345, BDSC51789); nosGal4 (BDSC64277); sax RNAi (TRiP.HMJ02118, BDSC42546); Mad RNAi (TRiP.JF01264, BDSC31316); UAS-Dpp (BDSC1486). For wildtype controls, *yw* (BDSC189) was used. UAS-brk.ORF.3xHA.GW (F000571) UAS-Mad.ORF.3xHA.GW (F001716) were obtained from FlyORF. UASGFP-αTubulin was gifts from Yukiko M. Yamashita.

### Immunofluorescence staining

Testes were dissected in PBS and fixed in 4% paraformaldehyde/PBS for 30 min. Samples were then washed in PBST (PBS + 0.2% TritonX-100) for 60 minutes, followed by overnight incubation with primary antibody in PBST containing 3% bovine serum albumin (BSA) at 4°C. Samples were then washed for 60 minutes in PBST, incubated with secondary antibody in 3% BSA in PBST at room temperature for 2 hours and then washed for 60 minutes (three times for 20 minutes each) in PBST. Samples were then mounted using VECTASHIELD with DAPI. The primary antibodies used were as follows: rat anti-Vasa (RRID: AB_760351, 1:20; DSHB); mouse anti-Hts

(1B1, 1:20; DSHB) mouse-anti-FasIII (1:20, 7G10; DSHB); Rabbit anti-HA C29F4 (RRID: AB_1549585, 1:300, Cell Signaling Technology, Cat# 3724); AlexaFluor-conjugated secondary antibodies (Abcam) were used at a 1:200 dilution. Images were acquired on a Zeiss LSM800 airyscan with a 63X/1.4 NA oil objective. Images were processed by image J/FIJI^42^.

### RNA in Situ Hybridization

For the Hybridization chain reaction method, we used a *brk* probe (kind gift from Leslie Dunipace and Angelike Stathopoulos) as performed as described previously^38^. Briefly, testes collected from 3-day-old flies were fixed in 1ml of 4% formaldehyde/PBS for 30 min, then permeabilized in phosphate-buffered saline (PBS) containing 1% Triton X-100 for 2 hours at room temperature, then washed with PBST (0.2% TritonX-100), 50/50 PBST/5xSSCT (0.1% Tween20) for 5 min, then 5xSSCT 5min at room temperature. Samples were prehybridized in probe hybridization buffer (Molecular Instruments) for 30 min at 37°C, followed by overnight hybridization (∼16 hours) with 0.8 pmol of probe at 37°C. After hybridization, specimens were equilibrated in amplification buffer (Molecular Instruments) for 5 min at room temperature. Hairpin solutions (Molecular Instruments) with 6 pmol of each hairpin were incubated for 90 seconds at 95°C, then cooled down in the dark at room temperature for 30 min. The cooled hairpins were added to the sample in amplification buffer, then incubated overnight (∼16 h) at room temperature. Next day, samples were washed with 5x SSC for 5min x2, 30min x1 and 5min x1, then followed by overnight incubation with primary antibody in PBST containing 3% bovine serum albumin (BSA) at 4°C. Samples were then washed for 60 minutes (20 minutes x3) in PBST (PBS + 0.2% TritonX-100), incubated with secondary antibody in 3% BSA in PBST at room temperature for 2 hours and then washed for 60 minutes (three times for 20 minutes each) in PBST. Samples were mounted using VECTASHIELD with 4’,6-diamidino-2-phenylindole (DAPI) (Vector Lab). Images were acquired on a Zeiss LSM800 airyscan with a 63X/1.4 NA oil objective.

For the single molecule FISH, Quasar 570 labeled probe against *bam* coding sequence was obtained from LGC Biosearch Technologies. For visualization of germ cells, NosGal4>αTubulin-GFP was used. Testes were dissected in 1X PBS then fixed in 1ml of 4% formaldehyde/PBS for 45 minutes. Fixed testes were rinsed 2 times with 1 ml of 1X PBS, then resuspended in 1ml of ice-cold 70% EtOH, and incubated for 1 hour-overnight at 4 °C. Testes were rinsed briefly in 1 ml of wash buffer (2X SSC and 10% deionized formamide), then incubated overnight at 37 °C for 16 hours in the dark with 50 nM of Stellaris probes in 200 μl of Hybridization Buffer containing 2X SSC, 10% dextran sulfate (MilliporeSigma), 1 μg/μl of yeast tRNA (MilliporeSigma), 2 mM vanadyl ribonucleoside complex (NEB), 0.02% RNase-free BSA (ThermoFisher), and 10% deionized formamide. Then, testes were washed 2 times for 30 minutes each at 37 °C in the dark in 1ml of prewarmed wash buffer (2X SSC, 10% formamide) and resuspended in a drop of VECTASHIELD with DAPI. Images were acquired on a Zeiss LSM800 airyscan with a 63X/1.4 NA oil objective.

### Quantification of Bam-mGL intensities

The generation of Bam-mGL or Bam(mut)-mGL reporter lines was previously described^22^. To assess the mGL intensity, short-term live imaging was used^7^. Testes from newly eclosed flies were dissected in Schneider’s Drosophila medium supplemented with 10% fetal bovine serum and glutamine–penicillin–streptomycin. Testes were placed on Gold Seal Rite-On Micro Slides with etched rings filled with media and covered with coverslips. Images were acquired on a Zeiss LSM800 airyscan with a 63X/1.4 NA oil objective. All images were taken using the same acquisition setting. The intensity of Bam-mGL or Bam(mut)-mGL was measured in the 4-, 8-, and 16-cell cyst SG regions. For intensity analysis, signals from each region were measured in ImageJ/Fiji, with background signals (from the same testis) subtracted.

### Statistical analysis and graphing

No statistical methods were used to predetermine sample size. All experiments were repeated at least two times to ensure reproducibility. The experiments were not randomized. The investigators were not blinded to allocation during experiments and outcome assessment. Statistical analysis and graphing were performed using GraphPad Prism 10 software. Data are means and standard deviations. All plotted data points are provided in Source Data. Individual numerical values displayed in all graphs are provided in Source data1.

## Supporting information

Sourcedata_1

## Acknowledgements

We would like to thank Amelie Raz, Leslie Dunipace and Stuart Newfeld for discussions and helpful suggestions; Leslie Dunipace, Angelike Stathopoulos, Yukiko Yamashita, and the Bloomington Drosophila Stock Center for reagents. This research is supported by R35GM128678 from the National Institute for General Medical Sciences and a start-up fund from UConn Health (to M.I.).

## Author contributions

M.I. and S.P, conceived the project, designed and executed experiments and analyzed data. J.G. participated in data analysis. All authors wrote and edited the manuscript.

---

**Figure S1.**
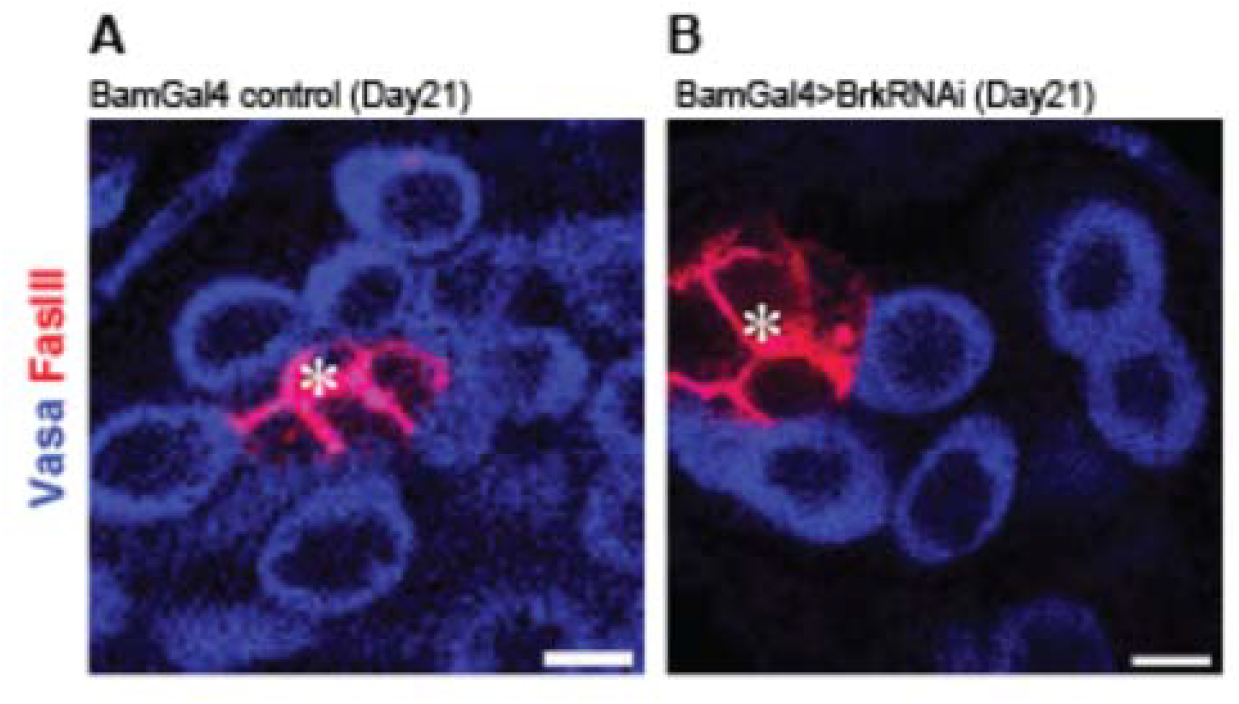
Brk is required for maintenance of stem cell pool. (A, B) Testis-tip images used for GSC number counting for indicated genotypes. The hub is visualized by FasIII (red) staining. Germ cells are visualized by Vasa staining (blue). GSCs were counted as germ cells directly attached to the hub. Imaging was performed to cover entire niche by using z-stacks of 1µm intervals. Asterisks indicate the hub. All scale bars indicate 10µm.

## References

1. Sheng, X.R., Brawley, C.M., and Matunis, E.L. (2009). Dedifferentiating Spermatogonia Outcompete Somatic Stem Cells for Niche Occupancy in the Drosophila Testis. Cell Stem Cell 5, 191–203. 10.1016/j.stem.2009.05.024.

2. Sheng, X.R., and Matunis, E. (2011). Live imaging of the Drosophila spermatogonial stem cell niche reveals novel mechanisms regulating germline stem cell output. Development 138, 3367–3376. 10.1242/dev.065797.

3. Herrera, S.C., and Bach, E.A. (2018). JNK signaling triggers spermatogonial dedifferentiation during chronic stress to maintain the germline stem cell pool in the Drosophila testis. eLife 7, e36095. 10.7554/eLife.36095.

4. Cheng, J., Türkel, N., Hemati, N., Fuller, M.T., Hunt, A.J., and Yamashita, Y.M. (2008). Centrosome misorientation reduces stem cell division during ageing. Nature 456, 599–604. 10.1038/nature07386.

5. Brawley, C., and Matunis, E. (2004). Regeneration of male germline stem cells by spermatogonial dedifferentiation in vivo. Science 304, 1331–1334. 10.1126/science.1097676.

6. Wallenfang, M.R., Nayak, R., and DiNardo, S. (2006). Dynamics of the male germline stem cell population during aging of Drosophila melanogaster. Aging Cell 5, 297–304. 10.1111/j.1474-9726.2006.00221.x.

7. Bener, M.B., Twillie, A., and Inaba, M. (2023). Dedifferentiating germ cells regain stem-cell specific polarity checkpoint prior to niche reentry. bioRxiv, 2023.2004.2026.538507. 10.1101/2023.04.26.538507.

8. Greenspan, L.J., and Matunis, E.L. (2017). Live Imaging of the Drosophila Testis Stem Cell Niche. Methods Mol Biol 1463, 63–74. 10.1007/978-1-4939-4017-2_4.

9. Greenspan, L.J., de Cuevas, M., and Matunis, E. (2015). Genetics of gonadal stem cell renewal. Annu Rev Cell Dev Biol 31, 291–315. 10.1146/annurev-cellbio-100913-013344.

10. Dansereau, D.A., and Lasko, P. (2008). The development of germline stem cells in Drosophila. Methods Mol Biol 450, 3–26. 10.1007/978-1-60327-214-8_1.

11. Yamashita, Y.M., Jones, D.L., and Fuller, M.T. (2003). Orientation of asymmetric stem cell division by the APC tumor suppressor and centrosome. Science 301, 1547–1550. 10.1126/science.1087795.

12. Yamashita, Y.M., Mahowald, A.P., Perlin, J.R., and Fuller, M.T. (2007). Asymmetric inheritance of mother versus daughter centrosome in stem cell division. Science 315, 518–521. 10.1126/science.1134910.

13. Inaba, M., Yuan, H., Salzmann, V., Fuller, M.T., and Yamashita, Y.M. (2010). E-cadherin is required for centrosome and spindle orientation in Drosophila male germline stem cells. PLoS One 5, e12473. 10.1371/journal.pone.0012473.

14. Inaba, M., Venkei, Z.G., and Yamashita, Y.M. (2015). The polarity protein Baz forms a platform for the centrosome orientation during asymmetric stem cell division in the Drosophila male germline. Elife 4, e04960.

15. Shivdasani, A.A., and Ingham, P.W. (2003). Regulation of stem cell maintenance and transit amplifying cell proliferation by tgf-beta signaling in Drosophila spermatogenesis. Curr Biol 13, 2065–2072. 10.1016/j.cub.2003.10.063.

16. Leatherman, J.L., and Dinardo, S. (2010). Germline self-renewal requires cyst stem cells and stat regulates niche adhesion in Drosophila testes. Nat Cell Biol 12, 806–811. 10.1038/ncb2086.

17. Tulina, N., and Matunis, E. (2001). Control of stem cell self-renewal in Drosophila spermatogenesis by JAK-STAT signaling. Science 294, 2546–2549. 10.1126/science.1066700.

18. Kawase, E., Wong, M.D., Ding, B.C., and Xie, T. (2004). Gbb/Bmp signaling is essential for maintaining germline stem cells and for repressing bam transcription in the Drosophilatestis. Development 131, 1365–1375. 10.1242/dev.01025.

19. Shivdasani, A.A., and Ingham, P.W. (2003). Regulation of Stem Cell Maintenance and Transit Amplifying Cell Proliferation by TGF-β Signaling in Drosophila Spermatogenesis. Current Biology 13, 2065–2072. 10.1016/j.cub.2003.10.063.

20. Schulz, C., Kiger, A.A., Tazuke, S.I., Yamashita, Y.M., Pantalena-Filho, L.C., Jones, D.L., Wood, C.G., and Fuller, M.T. (2004). A misexpression screen reveals effects of bag-of-marbles and TGF beta class signaling on the Drosophila male germ-line stem cell lineage. Genetics 167, 707–723. 10.1534/genetics.103.023184.

21. Beard, E.K., Norris, R.P., Furusho, M., Terasaki, M., and Inaba, M. (2025). Soma-to-germline BMP signal is essential for Drosophila spermiogenesis. Dev Biol 517, 140–147. 10.1016/j.ydbio.2024.09.016.

22. Ridwan, S.M., Twillie, A., Poursaeid, S., Beard, E.K., Bener, M.B., Antel, M., Cowan, A.E., Matsuda, S., and Inaba, M. (2024). Diffusible fraction of niche BMP ligand safeguards stem-cell differentiation. Nature Communications 15, 1166. 10.1038/s41467-024-45408-7.

23. Inaba, M., and Yamashita, Y.M. (2012). Asymmetric stem cell division: precision for robustness. Cell stem cell 11, 461–469.

24. Inaba, M., Buszczak, M., and Yamashita, Y.M. (2015). Nanotubes mediate niche-stem-cell signalling in the Drosophila testis. Nature 523, 329–332. 10.1038/nature14602.

25. Campbell, G., and Tomlinson, A. (1999). Transducing the Dpp morphogen gradient in the wing of Drosophila: regulation of Dpp targets by brinker. Cell 96, 553–562. 10.1016/s0092-8674(00)80659-5.

26. Jaźwińska, A., Kirov, N., Wieschaus, E., Roth, S., and Rushlow, C. (1999). The Drosophila gene brinker reveals a novel mechanism of Dpp target gene regulation. Cell 96, 563–573. 10.1016/s0092-8674(00)80660-1.

27. Minami, M., Kinoshita, N., Kamoshida, Y., Tanimoto, H., and Tabata, T. (1999). brinker is a target of Dpp in Drosophila that negatively regulates Dpp-dependent genes. Nature 398, 242–246. 10.1038/18451.

28. Saller, E., and Bienz, M. (2001). Direct competition between Brinker and Drosophila Mad in Dpp target gene transcription. EMBO Rep 2, 298–305. 10.1093/embo-reports/kve068.

29. Chen, Y., and Schüpbach, T. (2006). The role of brinker in eggshell patterning. Mechanisms of Development 123, 395–406. 10.1016/j.mod.2006.03.007.

30. Chen, D., and McKearin, D.M. (2003). A discrete transcriptional silencer in the bam gene determines asymmetric division of the Drosophila germline stem cell. Development 130, 1159–1170. 10.1242/dev.00325.

31. Chen, D., and McKearin, D. (2003). Dpp Signaling Silences bam Transcription Directly to Establish Asymmetric Divisions of Germline Stem Cells. Current Biology 13, 1786–1791. 10.1016/j.cub.2003.09.033.

32. Minami, M., Kinoshita, N., Kamoshida, Y., Tanimoto, H., and Tabata, T. (1999). brinker is a target of Dpp in Drosophila that negatively regulates Dpp-dependent genes. Nature 398, 242–246. 10.1038/18451.

33. Grant, C.E., Bailey, T.L., and Noble, W.S. (2011). FIMO: scanning for occurrences of a given motif. Bioinformatics 27, 1017–1018. 10.1093/bioinformatics/btr064.

34. Rauluseviciute, I., Riudavets-Puig, R., Blanc-Mathieu, R., Castro-Mondragon Jaime A., Ferenc, K., Kumar, V., Lemma, R.B., Lucas, J., Chèneby, J., Baranasic, D., et al. (2023). JASPAR 2024: 20th anniversary of the open-access database of transcription factor binding profiles. Nucleic Acids Research 52, D174–D182. 10.1093/nar/gkad1059.

35. Kirkpatrick, H., Johnson, K., and Laughon, A. (2001). Repression of dpp targets by binding of brinker to mad sites. J Biol Chem 276, 18216–18222. 10.1074/jbc.M101365200.

36. Antel, M., Raj, R., Masoud, M.Y.G., Pan, Z., Li, S., Mellone, B.G., and Inaba, M. (2022). Interchromosomal interaction of homologous Stat92E alleles regulates transcriptional switch during stem-cell differentiation. Nature Communications 13, 3981.

37. Sivasankaran, R., Vigano, M.A., Müller, B., Affolter, M., and Basler, K. (2000). Direct transcriptional control of the Dpp target omb by the DNA binding protein Brinker. Embo j 19, 6162–6172. 10.1093/emboj/19.22.6162.

38. Dunipace, L., Newcomb, S., and Stathopoulos, A. (2022). brinker levels regulated by a promoter proximal element support germ cell homeostasis. Development 149. 10.1242/dev.199890.

39. Insco, M.L., Leon, A., Tam, C.H., McKearin, D.M., and Fuller, M.T. (2009). Accumulation of a differentiation regulator specifies transit amplifying division number in an adult stem cell lineage. Proceedings of the National Academy of Sciences 106, 22311–22316. doi:10.1073/pnas.0912454106.

40. Bener, M.B., Slepchenko, B.M., and Inaba, M. (2025). Detection of dedifferentiated stem cells in Drosophila testis. bioRxiv. 10.1101/2025.03.06.641800.

41. Fuller, M.T., and Bate, M. (1993). The development of Drosophila melanogaster. Spermatogenesis 71, 147.

42. Schneider, C.A., Rasband, W.S., and Eliceiri, K.W. (2012). NIH Image to ImageJ: 25 years of image analysis. Nat Methods 9, 671–675. 10.1038/nmeth.2089.

